# White Matter Hyperintensity Distribution Differences in Aging and Neurodegenerative Disease Cohorts

**DOI:** 10.1101/2021.11.23.469690

**Authors:** Mahsa Dadar, Sawsan Mahmoud, Maryna Zhernovaia, Richard Camicioli, Josefina Maranzano, Simon Duchesne, For the CCNA Group

## Abstract

**Introduction:** White matter hyperintensities (WMHs) are common magnetic resonance imaging (MRI) findings in the aging population in general, as well as in patients with neurodegenerative diseases. They are known to exacerbate the cognitive deficits and worsen the clinical outcomes in the patients. However, it is not well-understood whether there are disease-specific differences in prevalence and distribution of WMHs in different neurodegenerative disorders.

**Methods:** Data included 976 participants with cross-sectional T1-weighted and fluid attenuated inversion recovery (FLAIR) MRIs from the Comprehensive Assessment of Neurodegeneration and Dementia (COMPASS-ND) cohort of the Canadian Consortium on Neurodegeneration in Aging (CCNA) with eleven distinct diagnostic groups: cognitively intact elderly (CIE), subjective cognitive impairment (SCI), mild cognitive impairment (MCI), vascular MCI (V-MCI), Alzheimer’s dementia (AD), vascular AD (V-AD), frontotemporal dementia (FTD), Lewy body dementia (LBD), cognitively intact elderly with Parkinson’s disease (PD-CIE), cognitively impaired Parkinson’s disease (PD-CI), and mixed dementias. WMHs were segmented using a previously validated automated technique. WMH volumes in each lobe and hemisphere were compared against matched CIE individuals, as well as each other, and between men and women.

**Results:** All cognitively impaired diagnostic groups had significantly greater overall WMH volumes than the CIE group. Vascular groups (i.e. V-MCI, V-AD, and mixed dementia) had significantly greater WMH volumes than all other groups, except for FTD, which also had significantly greater WMH volumes than all non-vascular groups. Women tended to have lower WMH burden than men in most groups and regions, controlling for age. The left frontal lobe tended to have a lower WMH burden than the right in all groups. In contrast, the right occipital lobe tended to have greater WMH loads than the left.

**Conclusions:** There were distinct differences in WMH prevalence and distribution across diagnostic groups, sexes, and in terms of asymmetry. WMH burden was significantly greater in all neurodegenerative dementia groups, likely encompassing areas exclusively impacted by neurodegeneration as well as areas related to cerebrovascular disease pathology.

## INTRODUCTION

White matter hyperintensities (WMHs) are areas of increased signal on T2-weighted and fluid attenuated inversion recovery (FLAIR) images that commonly occur in elderly individuals as well as patients with neurodegenerative diseases. WMHs have been associated with a multitude of underlying histopathological changes, such as myelin and axonal loss, loss of oligodendrocytes, microglial activation, lipohyalinosis, arteriosclerosis, vessel wall leakage, and collagen deposition in venular walls. Aetiologically, various pathophysiological mechanisms have been proposed, including ischemia, hypoperfusion, increased permeability of the blood–brain barrier, inflammation, degeneration and amyloid angiopathy (Gouw et al., 2010).

WMH burden has been linked to various risk factors in the elderly population without neurological comorbidities as well as in patients with neurodegenerative diseases. Age and hypertension are the main known risk factors of WMHs in the aging population in general, as well as hypercholesterolemia, diabetes, obesity, smoking, and alcoholism (Wardlaw et al., 2014; Morys et al., 2021; Grueter and Schulz, 2012). Similar associations with these risk factors have also been observed in MCI and AD populations (Dadar et al., 2020a). In contrast, in genetic frontotemporal dementia (FTD), severe WMH burden can be observed in absence of significant vascular risk factors and pathology, likely due to progranulin pathological processes (Caroppo et al., 2014; Woollacott et al., 2018). In Parkinson’s disease (PD), WMHs have also been linked to orthostatic hypotension and dysautonomia (Dadar et al., 2020b; Oh et al., 2013).

In terms of clinical symptoms, WMHs are associated with increased cognitive deficits (particularly in executive function and higher-order cognitive functions, such as planning, organizing, and monitoring behaviour) and neurobehavioral and psychiatric problems (e.g., depression, irritability, apathy, anxiety, and agitation), and gait difficulties (Anor et al., 2021; Arvanitakis et al., 2016; Baezner et al., 2008; Dadar et al., 2021, 2020a, 2020c; Kapasi et al., 2017; Misquitta et al., 2020; Srikanth et al., 2010; Teodorczuk et al., 2007; Van Der Flier et al., 2018). Of particular relevance, WMH are known to increase the risk of dementia for the same level of neurodegenerative pathology (Anor et al., 2021; Arvanitakis et al., 2016; Baezner et al., 2008; Dadar et al., 2021, 2020a, 2020c; Kapasi et al., 2017; Misquitta et al., 2020; Srikanth et al., 2010; Teodorczuk et al., 2007; Van Der Flier et al., 2018).

Previous studies in the literature have reported differences in WMH loads between patients with probable Alzheimer’s disease (AD), mild cognitive impairment (MCI), and age matched cognitively healthy controls (Barber et al., 1999; Dadar et al., 2018b; Desmarais et al., 2021; Tosto et al., 2014b). Similar results have been reported in patients with FTD and Lewy body dementia (LBD) (Barber et al., 1999; Desmarais et al., 2021; Anoop R. Varma et al., 2002). The findings have been more inconsistent in patients with PD, with some studies reporting significant differences between PD patients and matched controls, and others not (Dalaker et al., 2009b; Huang et al., 2020; Kandiah et al., 2014; Liu et al., 2021; Sunwoo et al., 2014). In general, later stage PD patients with MCI and dementia tend to have greater levels of WMH, whereas de novo cognitively normal PD patients seem to have similar WMH loads to controls (Butt et al., 2021; Dadar et al., 2018d; Liu et al., 2021), suggesting a possible link between WMHs and cognitive decline in PD.

Previous studies of neurodegenerative disorder have, in general, focussed on one disorder (e.g. pure AD without comorbid cerebrovascular pathology), limiting their ability to study the overlap between these disorders and recognize vascular contributions across neurodegenerative disorders. Investigating differences in prevalence and distribution of WMHs across different neurodegenerative diseases provides additional insights into the mechanisms through which cerebrovascular and neurodegenerative pathologies interact, possibly shedding light on potential synergistic relationships between theses pathologies.

Differences in image acquisition protocols and parameters, recruitment criteria, and variability in WMH assessment techniques has hindered the meaningful comparisons of WMH burden across different populations and studies. In essence, previous studies comparing WMH burden across different neurodegenerative diseases with the same image acquisition and WMH assessment methods have generally been performed in relatively small populations and with visual assessments of WMH burden, limiting their statistical power and sensitivity to detect the more subtle differences in WMH burden and distribution across cohorts (Burton et al., 2006; Erkinjuntti et al., 1994; Liu et al., 2021).

Our report addresses these issues. Taking advantage of the Comprehensive Assessment of Neurodegeneration and Dementia (COMPASS-ND) cohort of the Canadian Consortium on Neurodegeneration in Aging (CCNA) and our extensively validated automated WMH segmentation method developed to quantify WMHs in multi-center and multi-scanner studies of aging and neurodegeneration (Dadar et al., 2017b, 2017c), we have compared burden and distribution of WMHs across 11 diagnostic cohorts: cognitively intact elderly (CIE), subjective cognitive impairment (SCI), mild cognitive impairment (MCI), vascular MCI (V-MCI), Alzheimer’s dementia (AD), vascular AD (V-AD), frontotemporal dementia (FTD), Lewy body dementia (LBD), cognitively intact elderly with Parkinson’s disease (PD-CIE), cognitively impaired Parkinson’s disease (PD-CI), and mixed dementias as well as between men and women in each group. To our knowledge, this is the first study that consistently compares the distribution of WMHs across all these neurodegenerative disease populations using a large dataset (N=976) acquired consistently with a harmonized protocol.

## METHODS

### Participants

Data included 976 participants from the Comprehensive Assessment of Neurodegeneration and Dementia (COMPASS-ND) cohort of the CCNA, a national initiative to catalyze research on dementia (Chertkow et al., 2019). COMPASS-ND includes cognitively intact elderly subjects as well as patients with various forms of dementia and degenerative disorders and mild memory loss or subjective cognitive concerns. Ethical agreements were obtained at all respective sites. Written informed consent was obtained from all participants.

### Clinical diagnoses

Clinical diagnoses were determined by participating clinicians based on longitudinal clinical, screening, and MRI findings and based on standard diagnostic criteria (i.e. diagnosis reappraisal was performed using information from recruitment assessment, screening visit, clinical visit with physician input, and MRI). Diagnostic groups included, AD, CIE, FTD, LBD, MCI, PD-CIE, PD-MCI, PD-Dementia (for this study, PD-MCI and PD-Dementia groups were merged into one PD-CI group), SCI, V-AD, and V-MCI. For details on clinical group ascertainment, see Section 1 in the supplementary materials as well as <u>Pieruccini-Faria</u> et al. (Pieruccini-Faria et al., 2021).

### MRI data

All CCNA participants were scanned using the Canadian Dementia Imaging Protocol, a harmonized MRI protocol designed to reduce inter-scanner variability in multi-centric studies (Duchesne et al., 2019). The following sequences were used to detect WMHs:

- 3D isotropic T1w scans (voxel size = 1.0 × 1.0 × 1.0 mm^3^) with an acceleration factor of 2 (Siemens: MP-RAGE-PAT: 2; GE: IR-FSPGR-ASSET 1.5; Philips: TFE-Sense: 2)
- Fluid attenuated inversion recovery (T2w-FLAIR) images (voxel size = 0.9 × 0.9 × 3 mm^3^), fat saturation, and an acceleration factor of 2.

Table 1 shows the acquisition parameters for each sequence and scanner manufacturer. A detailed description, exam cards, and operators’ manual are publicly available at: www.cdip-pcid.ca.

**Table 1.**
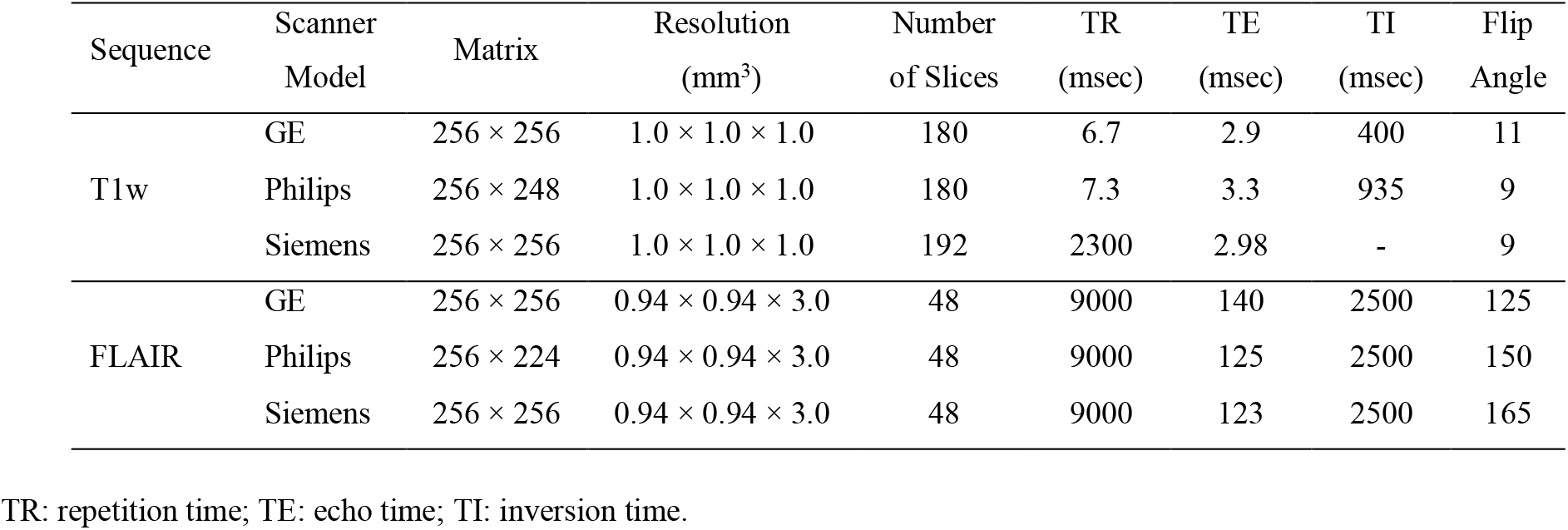
Acquisition parameters of the CDIP protocol.

### MRI preprocessing

All images were pre-processed using the following steps: image denoising (Coupe et al., 2008), intensity non-uniformity correction (Sled et al., 1998), and image intensity normalization into range 0-100. FLAIR images were co-registered (rigid registration, 6 parameters) to the T1w images with a mutual information cost function. The T1w scans were also linearly (Dadar et al., 2018a) and nonlinearly (Avants et al., 2008) registered to the MNI-ICBM152-2009c unbiased average template (Manera et al., 2020).

### WMH Segmentations

A previously validated segmentation method was used to segment WMHs (Dadar et al., 2017c, 2017b, 2018c). The technique employs a set of location and intensity features in combination with a random forests classifier to detect WMHs using T1w and FLAIR images. The training library consisted of manual, expert segmentations of WMHs from 60 participants from the CCNA (Dadar et al., 2017c, 2017b, 2018c). WMHs were automatically segmented for all participants in native FLAIR space.

#### Volumetric WMH Measures

The volumes of the WMHs for left and right frontal, parietal, temporal, and occipital lobes as well as the entire brain were calculated based on Hammers atlas (Dadar et al., 2018c; Hammers et al., 2003). The WMH volumes were normalized for intracranial volume to enable population comparisons and log-transformed to achieve normal distribution.

Figure 1 shows an example of the segmented WMHs and their separation into left and right frontal, parietal, temporal, and occipital lobes for one case.

**Figure 1.**
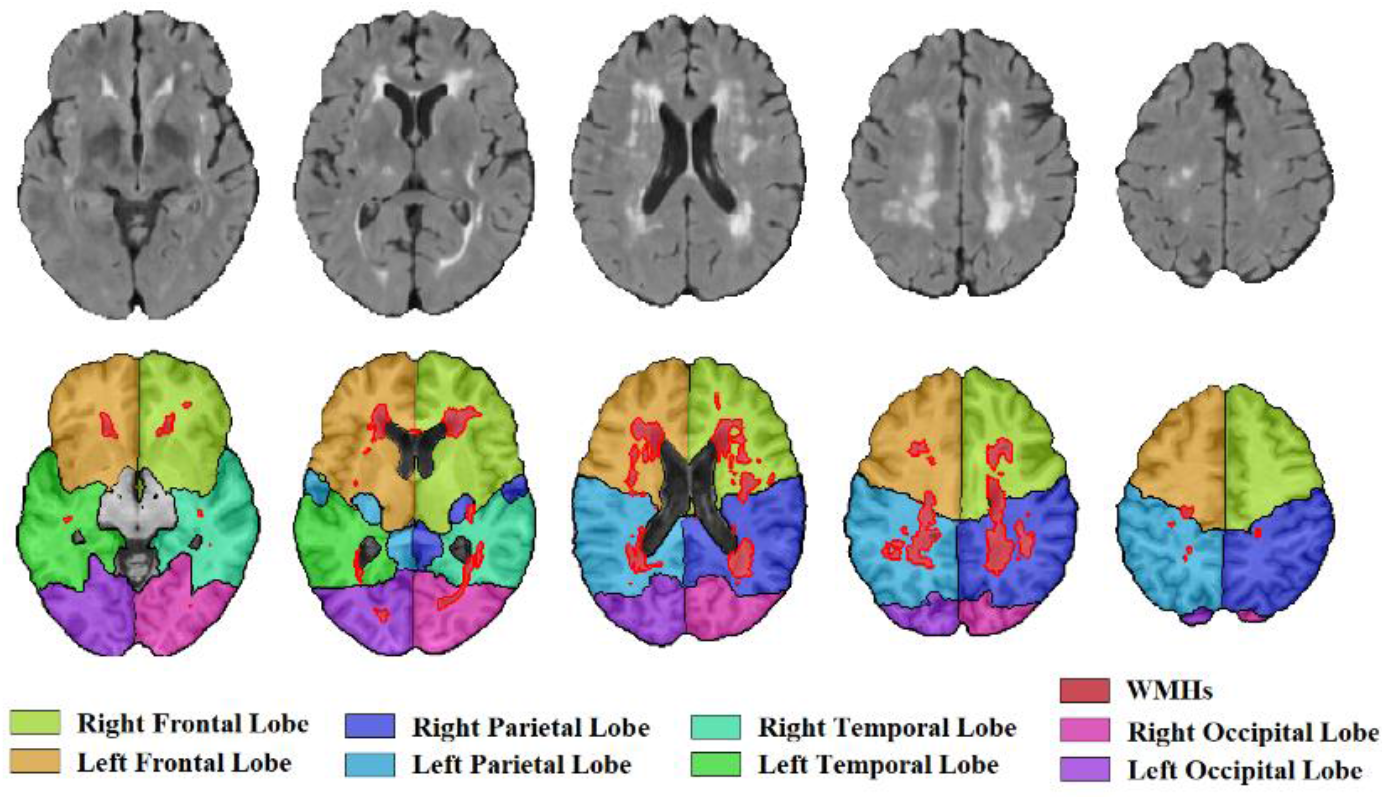
WMH segmentations and separation into left and right frontal, parietal, temporal, and occipital lobes.

#### WMH Distribution Maps

To obtain comparable voxel-wise WMH distribution maps, we nonlinearly registered all WMH masks to MNI-ICBM152-2009c template by concatenating linear FLAIR-to-T1w, linear T1w-to-ICBM, and nonlinear T1w-to-ICBM registration transformations.

### Statistical Analysis

The following linear regression models were used to assess differences between WMH volumes across different diagnoses and regions:

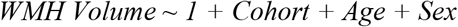

Where *WMH Volume* indicates total or lobar WMH burden and *Cohort* is the variable of interest indicating the contrast between each disease cohort and the reference (CIE) group. Similarly, pairwise regression models were run to assess differences between each diagnostic group pair.

The following regression models were used to assess sex differences in WMH burden in each group:

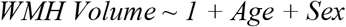

In these models, the variable of interest was *Sex*, contrasting women against men (self-reported sex at birth). The models were run separately for each diagnostic group. Paired t-tests were used to assess asymmetric differences between WMH volumes in the left and right hemispheres, contrasting the left hemisphere against the right. WMH volumes were normalized by lobe volumes in these analyses to ensure that slight differences in lobe volumes did not impact results.

WMH volumes were z-scored prior to the regression analyses. All results were corrected for multiple comparisons using false discovery rate (FDR) method with a significant threshold of 0.05. All statistical analyses were performed using MATLAB version 2021a.

### Data and Code Availability Statement

For information on availability and access to CCNA data, see http://ccna.dev.simalam.ca/compass-nd-study/. The WMH segmentation pipeline used is publicly available at http://nist.mni.mcgill.ca/white-matter-hyperintensities/.

## RESULTS

### MRI data

All preprocessed and registered images were visually assessed for quality control (presence of imaging artifacts, failure in registrations). WMH segmentations were also visually assessed for missing hyperintensities or over-segmentation. Either failure resulted in the participant being removed from the analyses (N = 9). All MRI processing, segmentation and quality control steps were blinded to clinical outcomes. Note that the provided data was already quality controlled by the CCNA imaging platform for presence of imaging artifacts, and only scans that had passed this quality control step were collected and used for this study. In final, 976 participants’ MRIs were entered in our analysis (see Table 2 for details).

**Table 2.** Demographic characteristics and WMH volumes (normalized values in cubic centimetres) for the participants used in this study.

| Diagnosis | N |  |  | Age | WMHs |  |  |  |  |
| --- | --- | --- | --- | --- | --- | --- | --- | --- | --- |
|  | Total | Female | Male |  | Total | Frontal | Parietal | Temporal | Occipital |
| AD | 88 | 37 | 51 | 74.54 ± 7.77 | 13.81 ± 9.58 | 6.13 ± 5.10 | 3.97 ± 3.75 | 1.69 ± 0.93 | 2.01 ± 1.15 |
| CIE | 105 | 78 | 27 | 69.99 ± 6.14 | 8.81 ± 6.57 | 4.14 ± 3.20 | 2.21 ± 2.35 | 1.20 ± 0.91 | 1.24 ± 0.88 |
| FTD | 37 | 20 | 17 | 65.65 ± 8.67 | 17.53 ± 12.57 | 8.06 ± 6.33 | 5.56 ± 5.14 | 1.81 ± 1.33 | 2.07 ± 1.33 |
| LBD | 26 | 3 | 23 | 72.49 ± 8.02 | 17.59 ± 13.52 | 7.00 ± 4.72 | 5.74 ± 5.73 | 2.11 ± 2.37 | 2.70 ± 1.93 |
| MCI | 264 | 112 | 152 | 72.09 ± 6.55 | 11.44 ± 10.01 | 5.34 ± 5.27 | 3.11 ± 3.96 | 1.40 ± 0.93 | 1.57 ± 1.08 |
| Mixed | 40 | 16 | 24 | 78.87 ± 6.45 | 35.28 ± 20.70 | 18.43 ± 11.13 | 10.55 ± 8.29 | 2.95 ± 1.90 | 3.31 ± 1.80 |
| PD-CIE | 76 | 35 | 41 | 67.05 ± 7.37 | 9.97 ± 9.41 | 4.90 ± 4.83 | 2.48 ± 3.56 | 1.24 ± 0.90 | 1.33 ± 0.96 |
| PD-CI | 51 | 7 | 44 | 72.70 ± 7.68 | 19.16 ± 18.52 | 8.99 ± 9.98 | 5.74 ± 6.86 | 2.14 ± 1.53 | 2.26 ± 1.48 |
| SCI | 132 | 98 | 34 | 70.55 ± 6.02 | 10.86 ± 9.21 | 5.63 ± 4.68 | 2.89 ± 3.73 | 1.25 ± 0.95 | 1.07 ± 3.85 |
| V-AD | 24 | 14 | 10 | 77.74 ± 7.17 | 28.55 ± 17.17 | 12.49 ± 10.29 | 9.40 ± 6.17 | 2.88 ± 1.29 | 3.72 ± 2.10 |
| V-MCI | 133 | 59 | 74 | 76.35 ± 6.27 | 29.21 ± 17.54 | 14.31 ± 9.58 | 9.34 ± 6.94 | 2.99 ± 1.75 | 2.53 ± 1.81 |

### WMH segmentations and volumes

The performance of the trained model was also evaluated against the manual expert labels in the training/validation subset (N = 60 participants), through 10-fold cross validation. Results accuracy was high (mean Dice similarity index = 0.80 ± 0.15).

Table 2 summarizes the WMH volumes (normalized values in cubic centimetres) in each diagnostic group.

Figure 2 shows the voxel-wise WMH distribution maps for each diagnostic group. WMHs were more prevalent in the periventricular regions for all groups, with the vascular groups (V-MCI, V-AD, and Mixed) having the highest WMH prevalence (see also Table 2). The FTD group also had high WMH prevalence, particularly considering that it included patients that were younger than all other diagnostic groups (mean age = 65.65).

**Figure 2.**
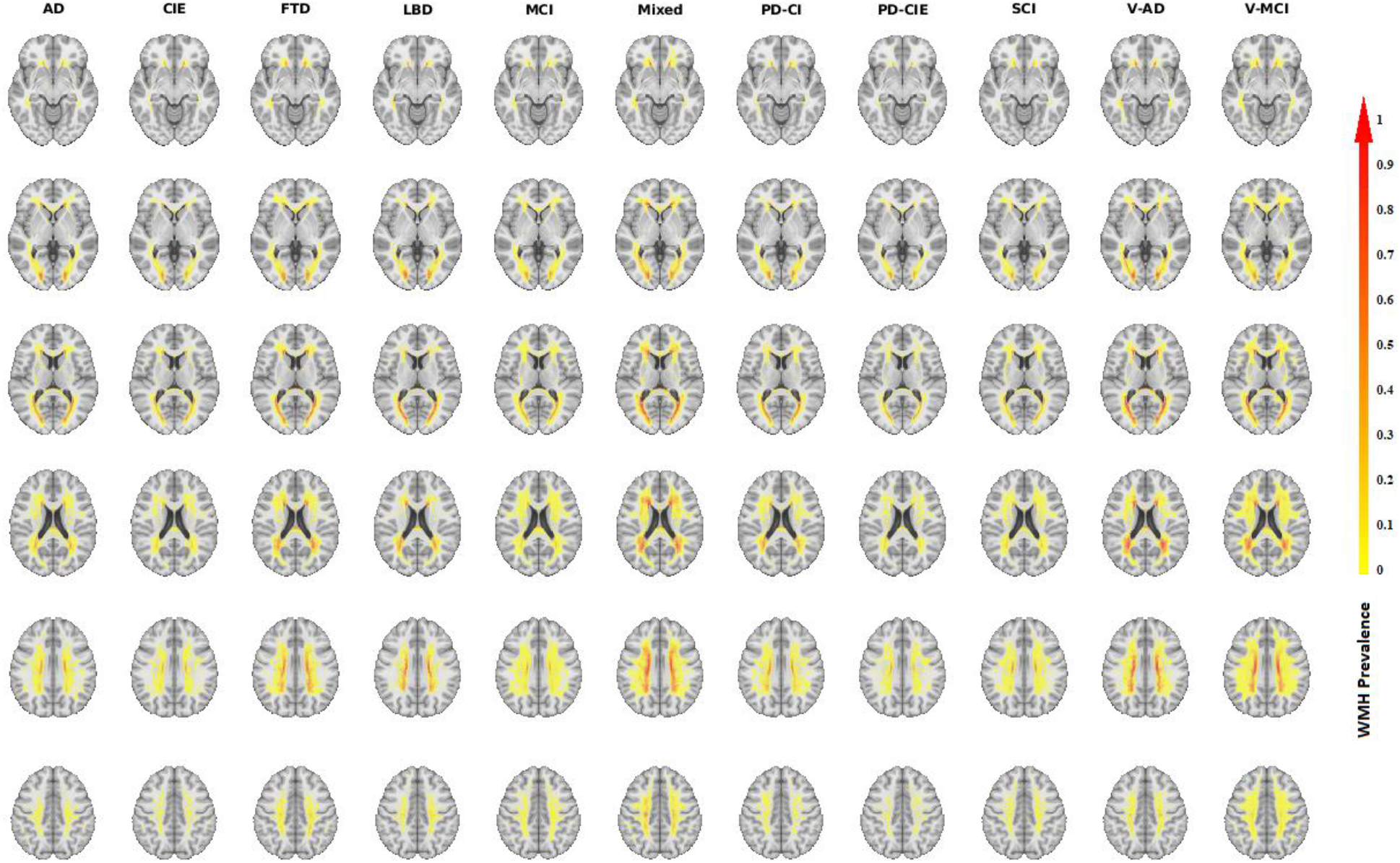
Voxel-wise WMH distribution maps for each diagnostic group.

### WMH volumes analyses

Table 3 shows the differences in total and regional WMH volumes for each group, contrasted against the CIE, and controlling for age and sex. The t statistic values contrast each diagnostic group versus CIE; i.e. a positive value indicates higher WMH in the diagnostic group than the CIE group. Figure 3 shows model β estimates for pairwise group differences for each diagnostic group pair and region. Significant differences after correction for multiple comparisons are marked with *.

**Table 3.** Total and regional WMH volume differences across diagnostic cohorts, controlling for age and sex. Values represent T statistics and uncorrected P values. Significant differences after FDR correction are shown in bold font.

| Region | Whole Brain | Left Frontal Lobe | Right Frontal Lobe | Left Parietal Lobe | Right Parietal Lobe | Left Temporal Lobe | Right Temporal Lobe | Left Occipital Lobe | Right Occipital Lobe |
| --- | --- | --- | --- | --- | --- | --- | --- | --- | --- |
| AD | **3.32, <0.001** | **2.26, 0.02** | 1.76, 0.07 | **2.15, 0.03** | **3.14, 0.002** | 1.43, 0.15 | **2.12, 0.03** | **2.16, 0.03** | **3.22, 0.001** |
| FTD | **7.15, <0.001** | **5.95, <0.001** | **6.34, <0.001** | **5.99, <0.001** | **6.80, <0.001** | **3.19, <0.001** | **4.44, <0.001** | **4.67, <0.001** | **3.69, <0.001** |
| LBD | **3.48, <0.001** | **2.52, 0.01** | **2.70, 0.007** | **3.56, <0.001** | **3.34, <0.001** | 1.26, 0.20 | 1.93, 0.05 | **3.30, <0.001** | **3.16, 0.002** |
| MCI | 1.53, 0.13 | 1.38, 0.16 | 1.14, 0.26 | 0.51, 0.61 | 1.07, 0.28 | 0.19, 0.85 | 0.25, 0.80 | -0.15, 0.88 | 1.35, 0.18 |
| Mixed | **9.35, <0.001** | **10.27, <0.001** | **9.44, <0.001** | **8.13, <0.001** | **7.55, <0.001** | **5.21, <0.001** | **5.06, <0.001** | **6.33, <0.001** | **4.71, <0.001** |
| PD-CI | **4.51, <0.001** | **4.41, <0.001** | **4.08, <0.001** | **4.18, <0.001** | **3.30, 0.001** | **2.99, 0.003** | **3.33, 0.001** | 1.99, 0.04 | **3.36, 0.001** |
| PD-CIE | 1.64, 0.10 | **2.42, 0.01** | 1.77, 0.08 | 1.47, 0.14 | 0.45, 0.65 | 0.52, 0.60 | 1.02, 0.30 | 0.33, 0.74 | 0.77, 0.44 |
| SCI | 2.04, 0.04 | **2.88, 0.004** | **2.70, 0.007** | 1.48, 0.14 | 1.29, 0.19 | 0.51, 0.60 | 0.04, 0.97 | -1.95, 0.05 | -0.99, 1.32 |
| V-AD | **6.82, <0.001** | **5.70, <0.001** | **5.23, <0.001** | **6.08, <0.001** | **6.83, <0.001** | **4.14, <0.001** | **5.55, <0.001** | **5.90, <0.001** | **6.60, <0.001** |
| V-MCI | **12.00, <0.001** | **12.07, <0.001** | **11.67, <0.001** | **10.62, <0.001** | **10.92, <0.001** | **8.43, <0.001** | **8.97, <0.001** | **4.19, <0.001** | **5.05, <0.001** |

**Figure 3.**
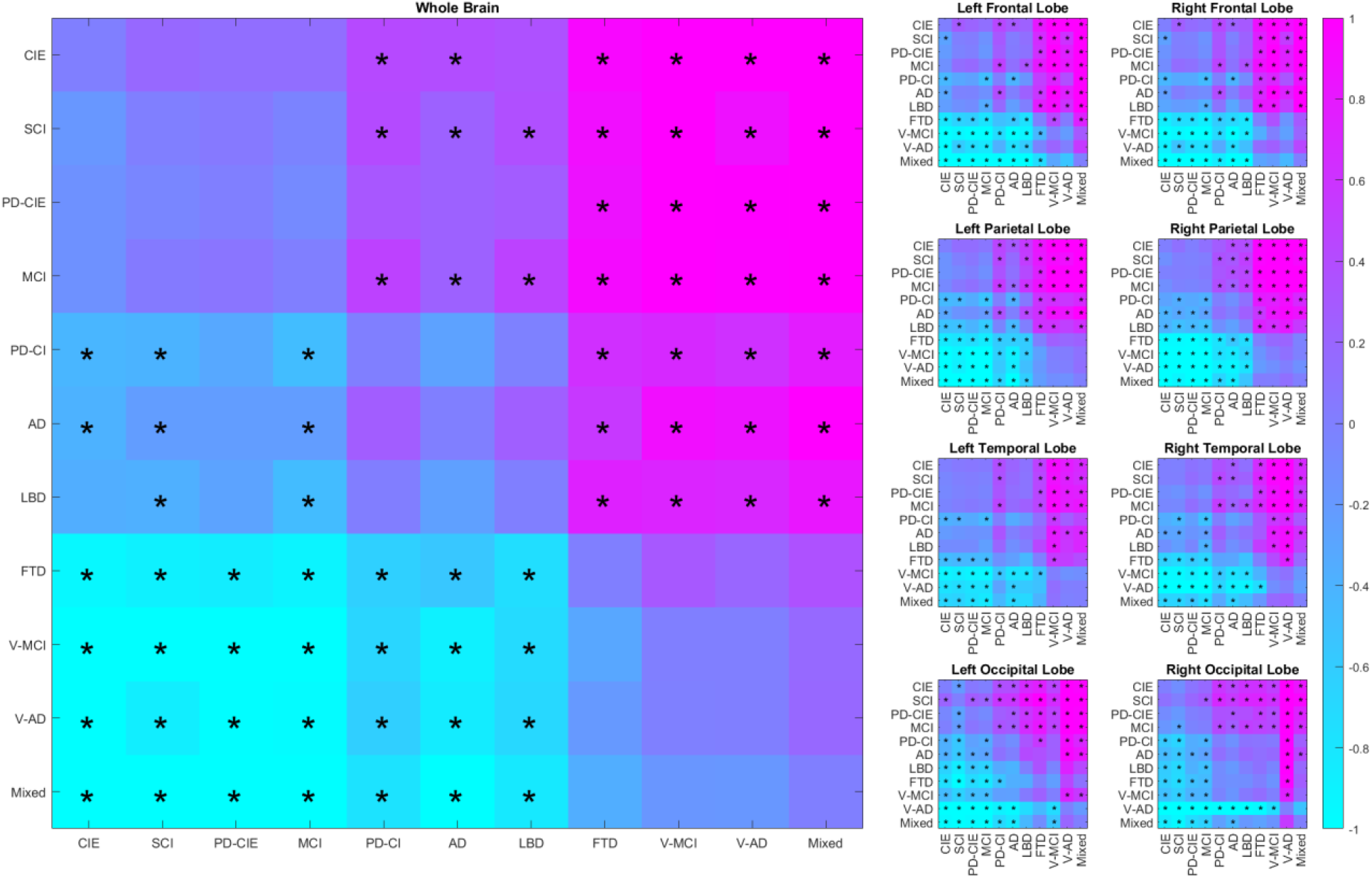
Regional and whole brain group differences in WMH volumes for each diagnostic pair. Colors indicate model β estimates. Significant differences after correction for multiple comparisons are marked with *.

The CIE group had significantly lower total WMH volumes than all cognitively impaired and dementia groups, except for the MCI group, likely since MCI participants with WMHs were classified in the vascular MCI (V-MCI) category. Participants with SCI had slightly higher total WMH volumes than the CIE group (uncorrected p value = 0.04), but this difference did not survive correction for multiple comparisons. As expected, V-MCI, V-AD, and Mixed groups had significantly higher WMH volumes than the CIE group in all regions. FTD group also had significantly greater WMH volumes than the CIE group in all regions. The (non-vascular) AD group also had significantly greater WMH loads than the CIE group overall and in left frontal, bilateral parietal and occipital, and right temporal lobes. The LBD group had significantly greater WMH volumes than the CIE group overall and in bilateral frontal, parietal, and occipital lobes. The cognitively impaired PD (PD-CI) group had significantly greater WMH loads than the CIE group overall and in all regions, except for left occipital lobe for which the significance did not survive correction for multiple comparisons. The PD-CIE group also had significantly greater WMH loads in left frontal lobe (uncorrected p value = 0.01), and marginally greater WMH loads in right frontal lobe (uncorrected p value = 0.08). Finally, the SCI group had significantly greater WMH loads than the CIE group in bilateral frontal lobes.

In the pairwise comparisons, vascular groups (i.e. V-MCI, V-AD, and Mixed) had significantly greater WMH volumes than all other groups, except for FTD, which also had significantly greater WMH volumes than all non-vascular groups.

Table 4 shows the differences in total and regional WMH volumes between men and women, controlling for age. The t statistic values contrast women versus men, i.e. a positive value indicates higher WMH in women than men, and vice versa. Overall, women tended to have lower WMH burden than men in most groups and regions, controlling for age. In the CIE group, women had significantly lower WMH volumes for whole brain WMHs, as well as right parietal and temporal lobes. In the FTD group, women had significantly lower WMH volumes for whole brain WMHs as well as right frontal lobe and left parietal and occipital lobes. In the MCI, V-MCI, and PD-CIE groups, women had significantly lower WMH volumes for bilateral occipital lobes.

**Table 4.**
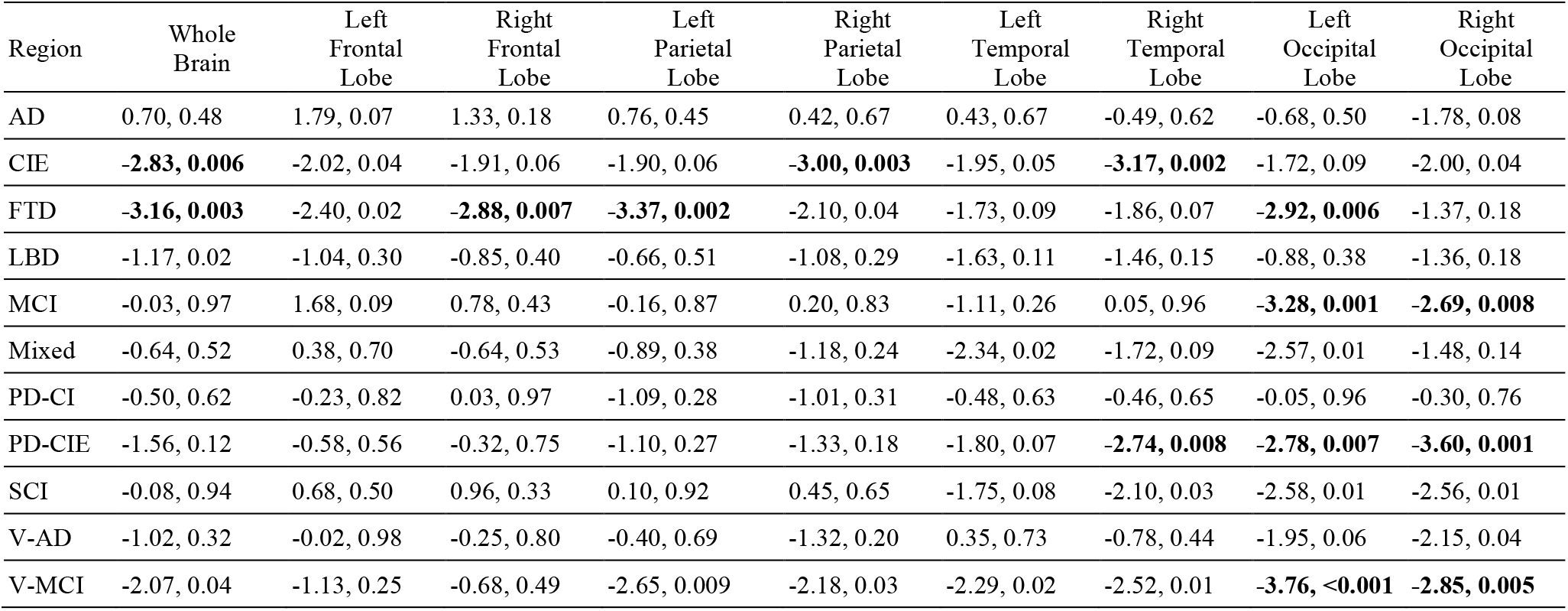
Total and regional WMH volume differences between men and women in each diagnostic cohort. Values represent T statistics and uncorrected P values. Significant differences after FDR correction are shown in bold font.

Table 5 shows the asymmetrical differences between left and right hemispheres (paired t-tests). The t statistic values contrast left versus right, i.e. a positive value indicates higher WMH in the left hemisphere than the right. Figure 4 shows boxplots of the normalized WMH values for each hemisphere, lobe, and diagnostic group. The left frontal lobe had lower WMH burden than the right in all groups, except for V-AD and mixed cohorts. In contrast, the right occipital lobe tended to have greater WMH loads than the left in all groups, except for PD-CI and V-AD cohorts. CIE, PD-CI and PD-CIE, and SCI groups had significantly higher WMH loads in the left parietal lobe, and AD, FTD, LBD, V-MCI, and V-AD groups had significantly greater WMH loads in the right temporal lobe. Figure 5 shows examples of significant asymmetry for each lobe. Figure S.1 shows the FLAIR images for the same subjects.

**Table 5.**
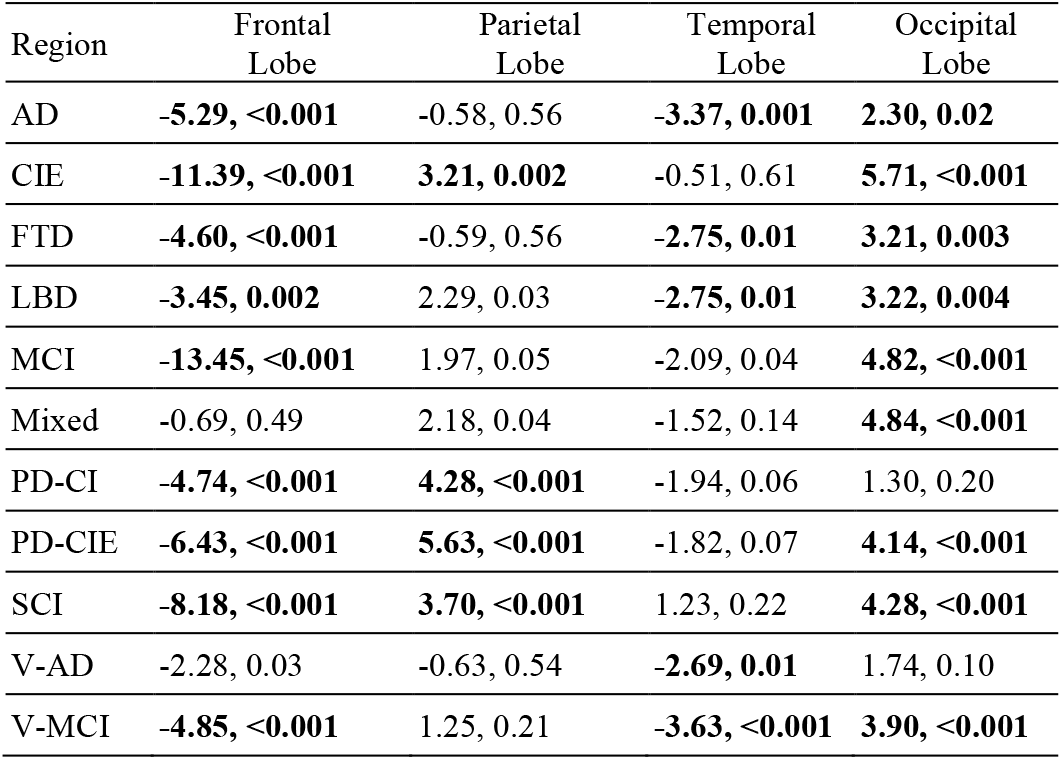
WMH volume differences between left and right hemispheres in each diagnostic cohort. Values represent T statistics and uncorrected P values. Significant differences after FDR correction are shown in bold font.

**Figure 4.**
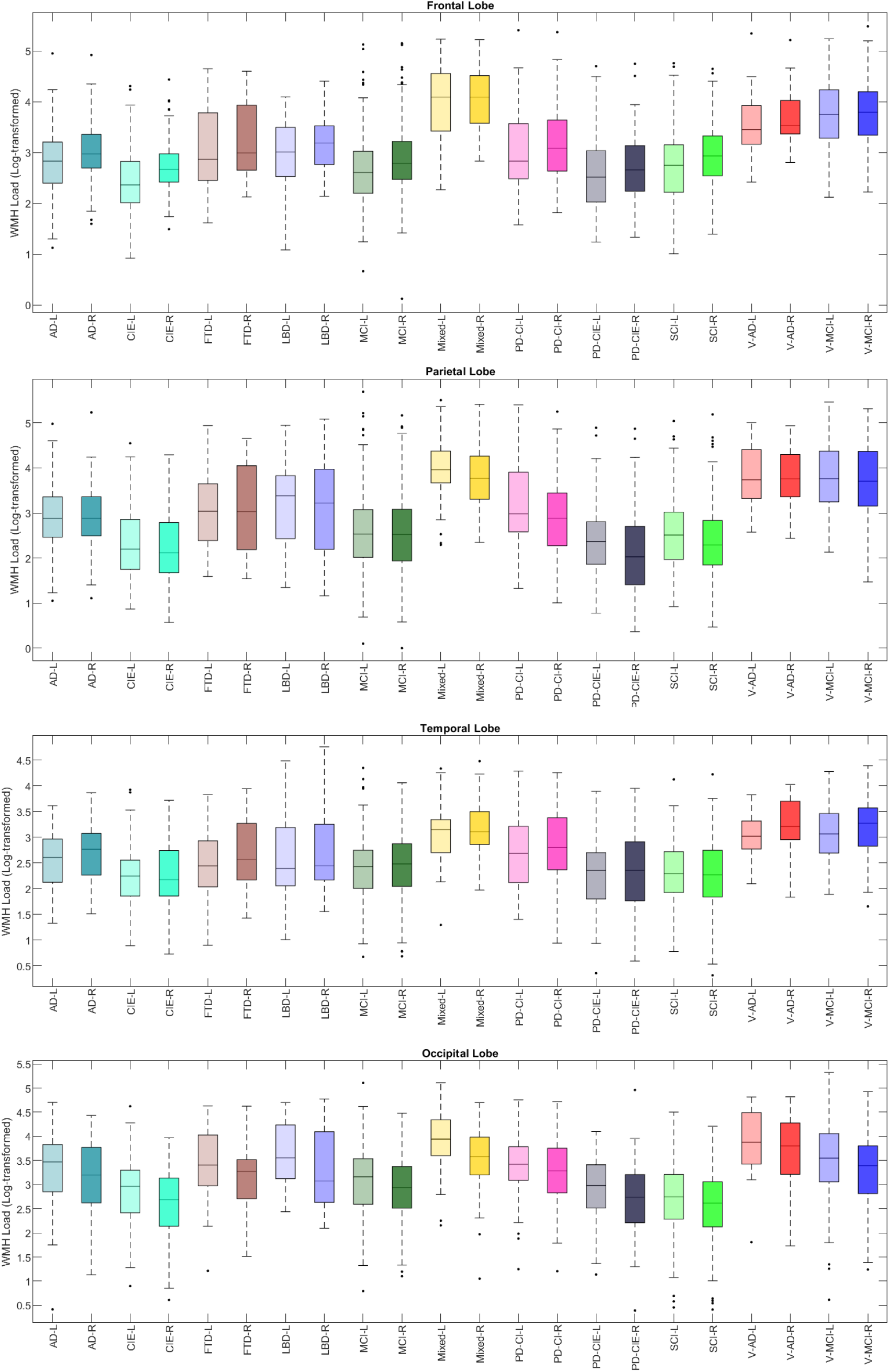
Boxplots of WMH volumes for each diagnostic group, lobe, and hemisphere. L= Left. R= Right.

**Figure 5.**
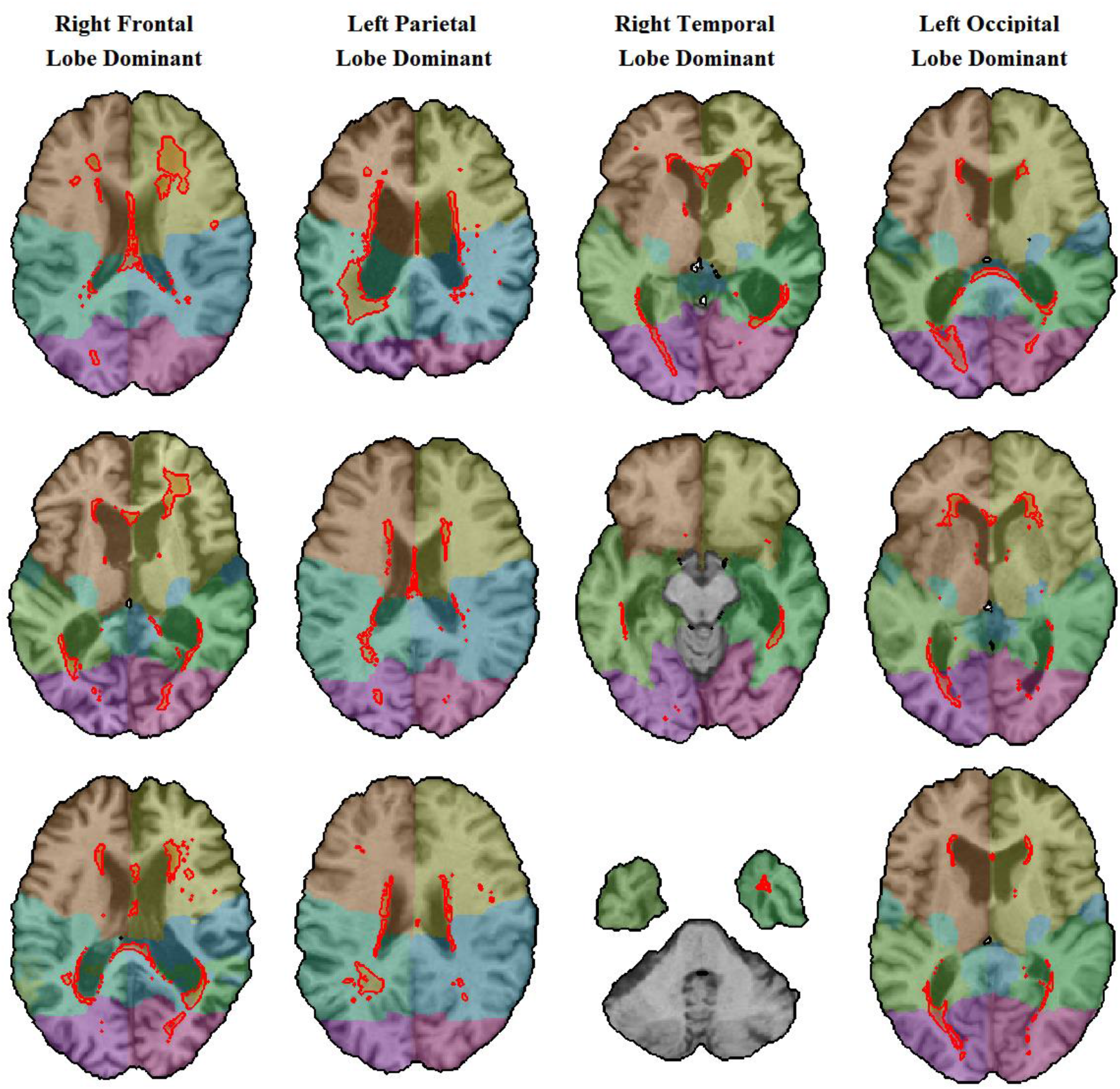
Example cases (3 of each) of right frontal (first column), left parietal (second column), right temporal (third column), and left occipital (fourth column) dominant asymmetry. The WMH segmentations are contoured in red.

## DISCUSSION

In this study, we compared the distribution of WMHs across eleven distinct neurodegenerative disease diagnostic groups using COMPASS-ND dataset from the CCNA acquired consistently with a harmonized protocol. Our results showed significantly greater WMH burden in all cognitively impaired and dementia groups compared with matched controls, whereas the cognitively intact PD (PD-CI) and SCI groups had slightly higher but non-significant differences.

As expected, the highest burden of WMHs was found in the mixed dementia group, followed by V-MCI and V-AD (Table 1). In patient groups that were not diagnosed with comorbid cerebrovascular pathology, the PD-CI, LBD, and FTD groups had the highest WMH loads. Interestingly, even the AD group not diagnosed with comorbid vascular pathology had significantly greater WMH burden than the controls, though this difference was much smaller (Tables 1 and 2). The higher burden of WMHs in MCI (Dadar et al., 2017a; DeCarli et al., 2001; Lopez et al., 2003) and AD patients (Burton et al., 2006; Capizzano et al., 2004; Dubois et al., 2014; Huynh et al., 2021; Tosto et al., 2014a) is relatively well established in the literature, whereas the results for other neurodegenerative disease cohorts have been more heterogeneous (Liu et al., 2021).

We found significantly greater overall and regional WMH loads in the PD-CI and LBD groups than the CIE, but not in PD-CIE. In addition, while the PD-CI group had higher WMH loads than the PD-CIE group (mean values of 19.16 cm^3^ versus 9.97 cm^3^ for PD-CI and PD-CIE, respectively, t stat = 1.9, uncorrected p value = 0.05), the difference did not survive correction for multiple comparisons (Figure 3). Similar to our findings, while studies in later stage PD patients generally report higher levels of WMHs (Liu et al., 2021; Mak et al., 2015; Piccini et al., 1995), studies investigating WMH differences in cognitively normal PD patients have not found significant differences between the patients and age matched controls (Dadar et al., 2018e; Dalaker et al., 2009a). Similarly, while some studies with smaller sample sizes have reported no differences in WMH burden between PD patients with dementia or LBD patients and healthy controls (Burton et al., 2006), others have found such significant differences (Barber et al., 1999), suggesting that larger cohorts are necessary to distinguish the more subtle differences (Butt et al., 2021).

Similar to other studies in the literature, FTD patients had significantly greater WMH burden than the CIE group (Desmarais et al., 2021; Huynh et al., 2021; A. R. Varma et al., 2002). In fact, the FTD group was the only non-vascular diagnostic group that had significantly greater overall WMH loads than all other groups, controlling for age and sex (Figure 3). This is particularly interesting, given that the FTD group was also the youngest group (Table 1), and older age is known to be the most significant correlate of WMHs. This might be in part due to the fact that high WMH burden can be observed in FTD patients in absence of significant vascular risk factors and pathology, possibly due to other pathological processes related to genetic mutations (Caroppo et al., 2014; Desmarais et al., 2021; Sudre et al., 2019; Woollacott et al., 2018). Unfortunately, our sample did not have genetic status information available, limiting our ability to investigate whether the FTD patients were carriers of specific mutations (e.g. <u>progranulin</u> gene), previously linked to increased WMH burden in genetic forms of FTD (Caroppo et al., 2014; Desmarais et al., 2021; Sudre et al., 2019; Woollacott et al., 2018).

The SCI group had significantly greater WMH loads in bilateral frontal lobes than the CIE (p < 0.007). While marginally higher (mean values of 10.86 cm^3^ versus 8.81 cm^3^ for SCI and CIE groups, respectively, t-stat = 2.04, uncorrected p value = 0.04), the overall WMH burden difference was not significant after correction for multiple comparisons. Previous reports on the association between WMHs and SCI have been varied. Van Rooden et al. reported significantly greater WMH volumes and Fazekas scores in SCI subjects compared to controls (van Rooden et al., 2018), while Caillund et al. did not find such differences (Caillaud et al., 2020). They did however report significant associations between WMHs and executive function in SCI (Caillaud et al., 2020). Similarly, Benedictus et al. linked presence of WMHs to future cognitive decline and progression to MCI or dementia (Benedictus et al., 2015).

While there are sex specific differences in causes and consequences of WMHs (Kumar and McCullough, 2021), the reports on sex differences in prevalence of WMHs have been inconsistent, with some studies not finding a significant difference (Wen et al., 2009; Ylikoski et al., 1995; Zhuang et al., 2018), some reporting greater WMH burden in women (Alqarni et al., 2021; Sachdev et al., 2009) and others reporting the opposite (Filomena et al., 2015; Geerlings et al., 2010). According to Simon et al., the greater WMH burden reported in women might be confounded by age, and be biased by premature death in men (which might be associated with cerebrovascular disease) (Simoni et al., 2012). In the present cohort, women tended to have lower WMH loads than men in general, controlling for age. In the CIE group, women had significantly lower WMH volumes for whole brain WMHs, as well as right parietal and temporal lobes. In the FTD group, women had significantly lower WMH volumes for whole brain WMHs as well as right frontal lobe and left parietal and occipital lobes. In the MCI, V-MCI, and PD-CIE groups, women had significantly lower WMH volumes for bilateral occipital lobes. Further investigations into associated vascular risk factors and co-morbidities are necessary to disentangle sex specific differences in prevalence of WMHs.

Regarding asymmetry, we found an overall greater burden of WMHs in right frontal and left occipital lobes (Figures 4 and 5). CIE, PD-CI and PD-CIE, and SCI groups had significantly higher WMH loads in the left parietal lobe, and AD, FTD, LBD, V-MCI, and V-AD groups had significantly greater WMH loads in the right temporal lobe. The presence of hemispheric dominance and predominance of WMH burden in right frontal lobe has been previously reported by Dhamoon et al. (Dhamoon et al., 2017). They also indicate that regional WMH volume asymmetry might be associated with lower function and functional decline (Dhamoon et al., 2017). Low et al. have also reported left-dominant occipital WMH burden in AD patients which was higher than that observed in MCI and control groups, as well as an association with poorer global cognition, memory, language, and executive functions among cognitively impaired participants (MCI and AD) (Low et al., 2019). There are established regional brain asymmetries in the frontal and occipital lobes which are also phylogenetically evident and have been detected in older primates (Toga and Thompson, 2003). These differences might also indicate differences in the evolutionary development of these lobes, leading to differences in susceptibility to neurodegeneration and presence of white matter pathology.

The image processing and WMH segmentation pipelines used in the present study have been developed and validated for use in multi-center and multi-scanner studies of aging and neurodegenerative disease populations and have been previously used in such applications (Anor et al., 2021; Dadar et al., 2020c, 2018e; Misquitta et al., 2020; Sanford et al., 2019). In addition to initial validation of the performance of the pipeline against gold standard manual segmentations which showed excellent agreement, manual quality control was performed to ensure the quality of the raw images, registrations, and segmentations.

Differences in image acquisition protocols, recruitment criteria, and WMH assessment techniques hinder reliable comparisons of WMH burden across different populations and studies. Using the COMPASS-ND cohort, acquired consistently with a harmonized protocol, we have compared burden and distribution of WMHs across 11 diagnostic cohorts, showing significantly greater WMH burden in all cognitively impaired and dementia groups compared with the matched controls, as well as asymmetric and sex specific trends. This emphasizes the need for further longitudinal investigations into the impact of WMHs in these neurodegenerative diseases as well as treatment and prevention strategies for vascular risk factors (anti-hypertensive medications, blood sugar management, lipid-lowering treatments, exercise, and lifestyle changes), which might slow down WMH progression and cognitive decline in neurodegenerative disease populations.

In these cohorts (even in cohorts not formally considered as mixed with vascular pathology), WMHs likely encompass areas exclusively impacted by neurodegeneration as well as areas related to non-specific MRI based small-vessel disease pathology. The cohorts formally classified as mixed, V-AD, and V-MCI, represent populations with more striking vascular pathogenic factors; hence the WMH volume of those groups are the highest among all cohorts. Future studies, refining and ranking the vascular pathogenic factors along with the MRI findings (e.g. presence of lacunar infarcts or macroscopic infarcts with clear vascular distribution patterns following vascular arterial territories) could help determine the extent of WMHs exclusively related to neurodegeneration versus small vessel disease and establish which component has a greater impact on cognitive decline and other clinical outcomes.

## Authors Contributions

**Mahsa Dadar**: Study concept and design, analysis of the data, drafting and revision of the manuscript.

**Sawsan Mahmoud:** Manual Segmentation of WMHs.

**Maryna Zhernovaia:** Manual Segmentation of WMHs.

**Richard Camicioli:** Study concept and design, interpretation of the data, revising the manuscript.

**Josefina Maranzano:** Study concept and design, interpretation of the data, revising the manuscript.

**Simon Duchesne:** Study concept and design, interpretation of the data, revising the manuscript.

## Acknowledgements

MD is supported by a scholarship from the Canadian Consortium on Neurodegeneration in Aging (CCNA) as well as by an Alzheimer Society Research Program (ASRP) postdoctoral award. CCNA is supported by a grant from the Canadian Institutes of Health Research with funding from several partners.

## References

Alqarni, A., Jiang, J., Crawford, J.D., Koch, F., Brodaty, H., Sachdev, P., Wen, W., 2021. Sex differences in risk factors for white matter hyperintensities in non-demented older individuals. Neurobiol. Aging 98, 197–204. https://doi.org/10.1016/j.neurobiolaging.2020.11.001

Anor, C.J., Dadar, M., Collins, D.L., Tartaglia, M.C., 2021. The Longitudinal Assessment of Neuropsychiatric Symptoms in Mild Cognitive Impairment and Alzheimer’s Disease and Their Association With White Matter Hyperintensities in the National Alzheimer’s Coordinating Center’s Uniform Data Set. Biol. Psychiatry Cogn. Neurosci. Neuroimaging, Imaging Biomarkers and Outcome Prediction 6, 70–78. https://doi.org/10.1016/j.bpsc.2020.03.006

Arvanitakis, Z., Capuano, A.W., Leurgans, S.E., Bennett, D.A., Schneider, J.A., 2016. Relation of cerebral vessel disease to Alzheimer’s disease dementia and cognitive function in elderly people: a cross-sectional study. Lancet Neurol. 15, 934–943.

Avants, B.B., Epstein, C.L., Grossman, M., Gee, J.C., 2008. Symmetric diffeomorphic image registration with cross-correlation: evaluating automated labeling of elderly and neurodegenerative brain. Med. Image Anal. 12, 26–41. https://doi.org/10.1016/j.media.2007.06.004

Baezner, H., Blahak, C., Poggesi, A., Pantoni, L., Inzitari, D., Chabriat, H., Erkinjuntti, T., Fazekas, F., Ferro, J.M., Langhorne, P., 2008. Association of gait and balance disorders with age-related white matter changes: the LADIS study. Neurology 70, 935–942.

Barber, R., Scheltens, P., Gholkar, A., Ballard, C., McKeith, I., Ince, P., Perry, R., O’brien, J., 1999. White matter lesions on magnetic resonance imaging in dementia with Lewy bodies, Alzheimer’s disease, vascular dementia, and normal aging. J. Neurol. Neurosurg. Psychiatry 67, 66–72.

Benedictus, M.R., van Harten, A.C., Leeuwis, A.E., Koene, T., Scheltens, P., Barkhof, F., Prins, N.D., van der Flier, W.M., 2015. White Matter Hyperintensities Relate to Clinical Progression in Subjective Cognitive Decline. Stroke 46, 2661–2664. https://doi.org/10.1161/STROKEAHA.115.009475

Burton, E.J., McKeith, I.G., Burn, D.J., Firbank, M.J., O’Brien, J.T., 2006. Progression of White Matter Hyperintensities in Alzheimer Disease, Dementia With Lewy Bodies, and Parkinson Disease Dementia: A Comparison With Normal Aging. Am. J. Geriatr. Psychiatry 14, 842–849. https://doi.org/10.1097/01.JGP.0000236596.56982.1c

Butt, A., Kamtchum-Tatuene, J., Khan, K., Shuaib, A., Jickling, G.C., Miyasaki, J.M., Smith, E.E., Camicioli, R., 2021. White matter hyperintensities in patients with Parkinson’s disease: A systematic review and meta-analysis. J. Neurol. Sci. 426, 117481. https://doi.org/10.1016/j.jns.2021.117481

Caillaud, M., Hudon, C., Boller, B., Brambati, S., Duchesne, S., Lorrain, D., Gagnon, J.-F., Maltezos, S., Mellah, S., Phillips, N., Consortium for the Early Identification of Alzheimer’s Disease-Quebec, Belleville, S., 2020. Evidence of a Relation Between Hippocampal Volume, White Matter Hyperintensities, and Cognition in Subjective Cognitive Decline and Mild Cognitive Impairment. J. Gerontol. Ser. B 75, 1382–1392. https://doi.org/10.1093/geronb/gbz120

Capizzano, A.A., Ación, L., Bekinschtein, T., Furman, M., Gomila, H., Martínez, A., Mizrahi, R., Starkstein, S.E., 2004. White matter hyperintensities are significantly associated with cortical atrophy in Alzheimer’s disease. J. Neurol. Neurosurg. Psychiatry 75, 822–827. https://doi.org/10.1136/jnnp.2003.019273

Caroppo, P., Le Ber, I., Camuzat, A., Clot, F., Naccache, L., Lamari, F., De Septenville, A., Bertrand, A., Belliard, S., Hannequin, D., Colliot, O., Brice, A., 2014. Extensive White Matter Involvement in Patients With Frontotemporal Lobar Degeneration: Think Progranulin. JAMA Neurol. 71, 1562–1566. https://doi.org/10.1001/jamaneurol.2014.1316

Chertkow, H., Borrie, M., Whitehead, V., Black, S.E., Feldman, H.H., Gauthier, S., Hogan, D.B., Masellis, M., McGilton, K., Rockwood, K., Tierney, M.C., Andrew, M., Hsiung, G.-Y.R., Camicioli, R., Smith, E.E., Fogarty, J., Lindsay, J., Best, S., Evans, A., Das, S., Mohaddes, Z., Pilon, R., Poirier, J., Phillips, N.A., MacNamara, E., Dixon, R.A., Duchesne, S., MacKenzie, I., Rylett, R.J., 2019. The Comprehensive Assessment of Neurodegeneration and Dementia: Canadian Cohort Study. Can. J. Neurol. Sci. 46, 499–511. https://doi.org/10.1017/cjn.2019.27

Coupe, P., Yger, P., Prima, S., Hellier, P., Kervrann, C., Barillot, C., 2008. An Optimized Blockwise Nonlocal Means Denoising Filter for 3-D Magnetic Resonance Images. IEEE Trans. Med. Imaging 27, 425–441. https://doi.org/10.1109/TMI.2007.906087

Dadar, M., Camicioli, R., Duchesne, S., Collins, D.L., 2020a. The temporal relationships between white matter hyperintensities, neurodegeneration, amyloid beta, and cognition. Alzheimers Dement. Diagn. Assess. Dis. Monit. 12, e12091. https://doi.org/10.1002/dad2.12091

Dadar, M., Fereshtehnejad, S.-M., Zeighami, Y., Dagher, A., Postuma, R.B., Collins, D.L., 2020b. White Matter Hyperintensities Mediate Impact of Dysautonomia on Cognition in Parkinson’s Disease. Mov. Disord. Clin. Pract. 7, 639–647. https://doi.org/10.1002/mdc3.13003

Dadar, M., Fonov, V.S., Collins, D.L., Initiative, A.D.N., 2018a. A comparison of publicly available linear MRI stereotaxic registration techniques. NeuroImage 174, 191–200.

Dadar, M., Gee, M., Shuaib, A., Duchesne, S., Camicioli, R., 2020c. Cognitive and motor correlates of grey and white matter pathology in Parkinson’s disease. NeuroImage Clin. 27, 102353. https://doi.org/10.1016/j.nicl.2020.102353

Dadar, M., Maranzano, J., Ducharme, S., Carmichael, O.T., Decarli, C., Collins, D.L., 2018b. Validation of T1w-based segmentations of white matter hyperintensity volumes in large-scale datasets of aging. Hum. Brain Mapp. 39, 1093–1107. https://doi.org/10.1002/hbm.23894

Dadar, M., Maranzano, J., Ducharme, S., Carmichael, O.T., Decarli, C., Collins, D.L., 2017a. Validation of T1w-based segmentations of white matter hyperintensity volumes in large-scale datasets of aging. Hum. Brain Mapp.

Dadar, M., Maranzano, J., Ducharme, S., Carmichael, O.T., Decarli, C., Collins, D.L., Initiative, A.D.N., 2018c. Validation of T 1w-based segmentations of white matter hyperintensity volumes in large-scale datasets of aging. Hum. Brain Mapp. 39, 1093–1107.

Dadar, M., Maranzano, J., Misquitta, K., Anor, C.J., Fonov, V.S., Tartaglia, M.C., Carmichael, O.T., Decarli, C., Collins, D.L., Alzheimer’s Disease Neuroimaging Initiative, 2017b. Performance comparison of 10 different classification techniques in segmenting white matter hyperintensities in aging. NeuroImage 157, 233–249. https://doi.org/10.1016/j.neuroimage.2017.06.009

Dadar, M., Miyasaki, J., Duchesne, S., Camicioli, R., 2021. White matter hyperintensities mediate the impact of amyloid ß on future freezing of gait in Parkinson’s disease. Parkinsonism Relat. Disord. 85, 95–101. https://doi.org/10.1016/j.parkreldis.2021.02.031

Dadar, M., Pascoal, T., Manitsirikul, S., Misquitta, K., Tartaglia, C., Brietner, J., Rosa-Neto, P., Carmichael, O., DeCarli, C., Collins, D.L., 2017c. Validation of a Regression Technique for Segmentation of White Matter Hyperintensities in Alzheimer’s Disease. IEEE Trans. Med. Imaging.

Dadar, M., Zeighami, Y., Yau, Y., Fereshtehnejad, S.-M., Maranzano, J., Postuma, R.B., Dagher, A., Collins, D.L., 2018d. White matter hyperintensities are linked to future cognitive decline in de novo Parkinson’s disease patients. NeuroImage Clin. 20, 892–900.

Dadar, M., Zeighami, Y., Yau, Y., Fereshtehnejad, S.-M., Maranzano, J., Postuma, R.B., Dagher, A., Collins, D.L., 2018e. White matter hyperintensities are linked to future cognitive decline in de novo Parkinson’s disease patients. NeuroImage Clin. 20, 892–900.

Dalaker, T.O., Larsen, J.P., Bergsland, N., Beyer, M.K., Alves, G., Dwyer, M.G., Tysnes, O.-B., Benedict, R.H., Kelemen, A., Bronnick, K., 2009a. Brain atrophy and white matter hyperintensities in early Parkinson’s disease. Mov. Disord. Off. J. Mov. Disord. Soc. 24, 2233–2241.

Dalaker, T.O., Larsen, J.P., Dwyer, M.G., Aarsland, D., Beyer, M.K., Alves, G., Bronnick, K., Tysnes, O.-B., Zivadinov, R., 2009b. White matter hyperintensities do not impact cognitive function in patients with newly diagnosed Parkinson’s disease. Neuroimage 47, 2083–2089.

DeCarli, C., Miller, B.L., Swan, G.E., Reed, T., Wolf, P.A., Carmelli, D., 2001. Cerebrovascular and brain morphologic correlates of mild cognitive impairment in the National Heart, Lung, and Blood Institute Twin Study. Arch. Neurol. 58, 643–647.

Desmarais, P., Gao, A.F., Lanctôt, K., Rogaeva, E., Ramirez, J., Herrmann, N., Stuss, D.T., Black, S.E., Keith, J., Masellis, M., 2021. White matter hyperintensities in autopsy-confirmed frontotemporal lobar degeneration and Alzheimer’s disease. Alzheimers Res. Ther. 13, 1–16.

Dhamoon, M.S., Cheung, Y.-K., Bagci, A., Alperin, N., Sacco, R.L., Elkind, M.S., Wright, C.B., 2017. Differential effect of left vs. right white matter hyperintensity burden on functional decline: the Northern Manhattan Study. Front. Aging Neurosci. 9, 305.

Dubois, B., Feldman, H.H., Jacova, C., Hampel, H., Molinuevo, J.L., Blennow, K., DeKosky, S.T., Gauthier, S., Selkoe, D., Bateman, R., others, 2014. Advancing research diagnostic criteria for Alzheimer’s disease: the IWG-2 criteria. Lancet Neurol. 13, 614–629.

Duchesne, S., Chouinard, I., Potvin, O., Fonov, V.S., Khademi, A., Bartha, R., Bellec, P., Collins, D.L., Descoteaux, M., Hoge, R., McCreary, C.R., Ramirez, J., Scott, C.J.M., Smith, E.E., Strother, S.C., Black, S.E., 2019. The Canadian Dementia Imaging Protocol: Harmonizing National Cohorts. J. Magn. Reson. Imaging 49, 456–465. https://doi.org/10.1002/jmri.26197

Erkinjuntti, T., Gao, F., Lee, D.H., Eliasziw, M., Merskey, H., Hachinski, V.C., 1994. Lack of Difference in Brain Hyperintensities Between Patients With Early Alzheimer’s Disease and Control Subjects. Arch. Neurol. 51, 260–268. https://doi.org/10.1001/archneur.1994.00540150054016

Filomena, J., Riba-Llena, I., Vinyoles, E., Tovar, J.L., Mundet, X., Castañé, X., Vilar, A., López-Rueda, A., Jiménez-Baladó, J., Cartanyà, A., Montaner, J., Delgado, P., 2015. Short-Term Blood Pressure Variability Relates to the Presence of Subclinical Brain Small Vessel Disease in Primary Hypertension. Hypertension 66, 634–640. https://doi.org/10.1161/HYPERTENSIONAHA.115.05440

Geerlings, M.I., Appelman, A.P.A., Vincken, K.L., Algra, A., Witkamp, T.D., Mali, W.P.T.M., van der Graaf, Y., 2010. Brain volumes and cerebrovascular lesions on MRI in patients with atherosclerotic disease. The SMART-MR study. Atherosclerosis 210, 130–136. https://doi.org/10.1016/j.atherosclerosis.2009.10.039

Gouw, A.A., Seewann, A., Van Der Flier, W.M., Barkhof, F., Rozemuller, A.M., Scheltens, P., Geurts, J.J., 2010. Heterogeneity of small vessel disease: a systematic review of MRI and histopathology correlations. J. Neurol. Neurosurg. Psychiatry jnnp-2009.

Grueter, B.E., Schulz, U.G., 2012. Age-related cerebral white matter disease (leukoaraiosis): a review. Postgrad. Med. J. 88, 79–87. https://doi.org/10.1136/postgradmedj-2011-130307

Hammers, A., Allom, R., Koepp, M.J., Free, S.L., Myers, R., Lemieux, L., Mitchell, T.N., Brooks, D.J., Duncan, J.S., 2003. Three-dimensional maximum probability atlas of the human brain, with particular reference to the temporal lobe. Hum. Brain Mapp. 19, 224–247.

Huang, X., Wen, M.-C., Ng, S.Y.-E., Hartono, S., Chia, N.S.-Y., Choi, X., Tay, K.-Y., Au, W.-L., Chan, L.-L., Tan, E.-K., Tan, L.C.-S., 2020. Periventricular white matter hyperintensity burden and cognitive impairment in early Parkinson’s disease. Eur. J. Neurol. 27, 959–966. https://doi.org/10.1111/ene.14192

Huynh, K., Piguet, O., Kwok, J., Dobson-Stone, C., Halliday, G.M., Hodges, J.R., Landin-Romero, R., 2021. Clinical and Biological Correlates of White Matter Hyperintensities in Patients With Behavioral-Variant Frontotemporal Dementia and Alzheimer Disease. Neurology 96, e1743–e1754. https://doi.org/10.1212/WNL.0000000000011638

Kandiah, N., Zainal, N.H., Narasimhalu, K., Chander, R.J., Ng, A., Mak, E., Au, W.L., Sitoh, Y.Y., Nadkarni, N., Tan, L.C., 2014. Hippocampal volume and white matter disease in the prediction of dementia in Parkinson’s disease. Parkinsonism Relat. Disord. 20, 1203–1208.

Kapasi, A., DeCarli, C., Schneider, J.A., 2017. Impact of multiple pathologies on the threshold for clinically overt dementia. Acta Neuropathol. (Berl.) 134, 171–186.

Kumar, A., McCullough, L., 2021. Cerebrovascular disease in women. Ther. Adv. Neurol. Disord. 14, 1756286420985237. https://doi.org/10.1177/1756286420985237

Liu, H., Deng, B., Xie, F., Yang, X., Xie, Z., Chen, Y., Yang, Z., Huang, X., Zhu, S., Wang, Q., 2021. The influence of white matter hyperintensity on cognitive impairment in Parkinson’s disease. Ann. Clin. Transl. Neurol.

Lopez, O.L., Jagust, W.J., DeKosky, S.T., Becker, J.T., Fitzpatrick, A., Dulberg, C., Breitner, J., Lyketsos, C., Jones, B., Kawas, C., others, 2003. Prevalence and classification of mild cognitive impairment in the Cardiovascular Health Study Cognition Study: part 1. Arch. Neurol. 60, 1385–1389.

Low, A., Ng, K.P., Chander, R.J., Wong, B., Kandiah, N., 2019. Association of Asymmetrical White Matter Hyperintensities and Apolipoprotein E4 on Cognitive Impairment. J. Alzheimers Dis. 70, 953–964. https://doi.org/10.3233/JAD-190159

Mak, E., Dwyer, M.G., Ramasamy, D.P., Au, W.L., Tan, L., Zivadinov, R., Kandiah, N., 2015. White matter hyperintensities and mild cognitive impairment in Parkinson’s disease. J. Neuroimaging 25, 754–760.

Manera, A.L., Dadar, M., Fonov, V., Collins, D.L., 2020. CerebrA, registration and manual label correction of Mindboggle-101 atlas for MNI-ICBM152 template. Sci. Data 7, 1–9.

Misquitta, K., Dadar, M., Louis Collins, D., Tartaglia, M.C., 2020. White matter hyperintensities and neuropsychiatric symptoms in mild cognitive impairment and Alzheimer’s disease. NeuroImage Clin. 28, 102367. https://doi.org/10.1016/j.nicl.2020.102367

Morys, F., Dadar, M., Dagher, A., 2021. Association between mid-life obesity, its metabolic consequences, cerebrovascular disease and cognitive decline. J. Clin. Endocrinol. Metab. https://doi.org/10.1210/clinem/dgab135

Oh, Y.-S., Kim, J.-S., Lee, K.-S., 2013. Orthostatic and supine blood pressures are associated with white matter hyperintensities in Parkinson disease. J. Mov. Disord. 6, 23.

Piccini, P., Pavese, N., Canapicchi, R., Paoli, C., Dotto, P.D., Puglioli, M., Rossi, G., Bonuccelli, U., 1995. White Matter Hyperintensities in Parkinson’s Disease: Clinical Correlations. Arch. Neurol. 52, 191–194. https://doi.org/10.1001/archneur.1995.00540260097023

Pieruccini-Faria, F., Black, S.E., Masellis, M., Smith, E.E., Almeida, Q.J., Li, K.Z.H., Bherer, L., Camicioli, R., Montero-Odasso, M., 2021. Gait variability across neurodegenerative and cognitive disorders: Results from the Canadian Consortium of Neurodegeneration in Aging (CCNA) and the Gait and Brain Study. Alzheimers Dement. n/a. https://doi.org/10.1002/alz.12298

Sachdev, P.S., Parslow, R., Wen, W., Anstey, K.J., Easteal, S., 2009. Sex differences in the causes and consequences of white matter hyperintensities. Neurobiol. Aging 30, 946–956. https://doi.org/10.1016/j.neurobiolaging.2007.08.023

Sanford, R., Strain, J., Dadar, M., Maranzano, J., Bonnet, A., Mayo, N.E., Scott, S.C., Fellows, L.K., Ances, B.M., Collins, D.L., 2019. HIV infection and cerebral small vessel disease are independently associated with brain atrophy and cognitive impairment. AIDS Lond. Engl. 33, 1197–1205. https://doi.org/10.1097/QAD.0000000000002193

Simoni, M., Li, L., Paul, N.L.M., Gruter, B.E., Schulz, U.G., Küker, W., Rothwell, P.M., 2012. Age- and sex-specific rates of leukoaraiosis in TIA and stroke patients: Population-based study. Neurology 79, 1215–1222. https://doi.org/10.1212/WNL.0b013e31826b951e

Sled, J.G., Zijdenbos, A.P., Evans, A.C., 1998. A nonparametric method for automatic correction of intensity nonuniformity in MRI data. IEEE Trans. Med. Imaging 17, 87–97.

Srikanth, V., Phan, T.G., Chen, J., Beare, R., Stapleton, J.M., Reutens, D.C., 2010. The location of white matter lesions and gait—A voxel-based study. Ann. Neurol. Off. J. Am. Neurol. Assoc. Child Neurol. Soc. 67, 265–269.

Sudre, C.H., Bocchetta, M., Heller, C., Convery, R., Neason, M., Moore, K.M., Cash, D.M., Thomas, D.L., Woollacott, I.O.C., Foiani, M., Heslegrave, A., Shafei, R., Greaves, C., van Swieten, J., Moreno, F., Sanchez-Valle, R., Borroni, B., Laforce, R., Masellis, M., Tartaglia, M.C., Graff, C., Galimberti, D., Rowe, J.B., Finger, E., Synofzik, M., Vandenberghe, R., de Mendonça, A., Tagliavini, F., Santana, I., Ducharme, S., Butler, C., Gerhard, A., Levin, J., Danek, A., Frisoni, G.B., Sorbi, S., Otto, M., Zetterberg, H., Ourselin, S., Cardoso, M.J., Rohrer, J.D., Rossor, M.N., Warren, J.D., Fox, N.C., Guerreiro, R., Bras, J., Thomas, D.L., Nicholas, J., Mead, S., Jiskoot, L., Meeter, L., Panman, J., Papma, J., van Minkelen, R., Pijnenburg, Y., Barandiaran, M., Indakoetxea, B., Gabilondo, A., Tainta, M., Arriba, M. de, Gorostidi, A., Zulaica, M., Villanua, J., Diaz, Z., Borrego-Ecija, S., Olives, J., Lladó, A., Balasa, M., Antonell, A., Bargallo, N., Premi, E., Cosseddu, M., Gazzina, S., Padovani, A., Gasparotti, R., Archetti, S., Black, S., Mitchell, S., Rogaeva, E., Freedman, M., Keren, R., Tang-Wai, D., Öijerstedt, L., Andersson, C., Jelic, V., Thonberg, H., Arighi, A., Fenoglio, C., Scarpini, E., Fumagalli, G., Cope, T., Timberlake, C., Rittman, T., Shoesmith, C., Bartha, R., Rademakers, R., Wilke, C., Karnarth, H.-O., Bender, B., Bruffaerts, R., Vandamme, P., Vandenbulcke, M., Ferreira, C.B., Miltenberger, G., Maruta, C., Verdelho, A., Afonso, S., Taipa, R., Caroppo, P., Di Fede, G., Giaccone, G., Prioni, S., Redaelli, V., Rossi, G., Tiraboschi, P., Duro, D., Almeida, M.R., Castelo-Branco, M., Leitão, M.J., Tabuas-Pereira, M., Santiago, B., Gauthier, S., Rosa-Neto, P., Veldsman, M., Flanagan, T., Prix, C., Hoegen, T., Wlasich, E., Loosli, S., Schonecker, S., Semler, E., Anderl-Straub, S., Benussi, L., Binetti, G., Ghidoni, R., Pievani, M., Lombardi, G., Nacmias, B., Ferrari, C., Bessi, V., 2019. White matter hyperintensities in progranulin-associated frontotemporal dementia: A longitudinal GENFI study. NeuroImage Clin. 24, 102077. https://doi.org/10.1016/j.nicl.2019.102077

Sunwoo, M.K., Jeon, S., Ham, J.H., Hong, J.Y., Lee, J.E., Lee, J.-M., Sohn, Y.H., Lee, P.H., 2014. The burden of white matter hyperintensities is a predictor of progressive mild cognitive impairment in patients with P arkinson’s disease. Eur. J. Neurol. 21, 922–e50.

Teodorczuk, A., O’Brien, J.T., Firbank, M.J., Pantoni, L., Poggesi, A., Erkinjuntti, T., Wallin, A., Wahlund, L.-O., Gouw, A., Waldemar, G., 2007. White matter changes and late-life depressive symptoms: longitudinal study. Br. J. Psychiatry 191, 212–217.

Toga, A.W., Thompson, P.M., 2003. Mapping brain asymmetry. Nat. Rev. Neurosci. 4, 37–48.

Tosto, G., Zimmerman, M.E., Carmichael, O.T., Brickman, A.M., 2014a. Predicting aggressive decline in mild cognitive impairment: the importance of white matter hyperintensities. JAMA Neurol. 71, 872–877.

Tosto, G., Zimmerman, M.E., Carmichael, O.T., Brickman, A.M., Initiative, A.D.N., 2014b. Predicting aggressive decline in mild cognitive impairment: the importance of white matter hyperintensities. JAMA Neurol. 71, 872–877.

Van Der Flier, W.M., Skoog, I., Schneider, J.A., Pantoni, L., Mok, V., Chen, C.L., Scheltens, P., 2018. Vascular cognitive impairment. Nat. Rev. Dis. Primer 4, 1–16.

van Rooden, S., van den Berg-Huysmans, A.A., Croll, P.H., Labadie, G., Hayes, J.M., Viviano, R., van der Grond, J., Rombouts, S.A.R.B., Damoiseaux, J.S., 2018. Subjective Cognitive Decline Is Associated with Greater White Matter Hyperintensity Volume. J. Alzheimers Dis. 66, 1283–1294. https://doi.org/10.3233/JAD-180285

Varma, Anoop R., Laitt, R., Lloyd, J.J., Carson, K.J., Snowden, J.S., Neary, D., Jackson, A., 2002. Diagnostic value of high signal abnormalities on T2 weighted MRI in the differentiation of Alzheimer’s, frontotemporal and vascular dementias. Acta Neurol. Scand. 105, 355–364.

Varma, A. R., Laitt, R., Lloyd, J.J., Carson, K.J., Snowden, J.S., Neary, D., Jackson, A., 2002. Diagnostic value of high signal abnormalities on T2 weighted MRI in the differentiation of Alzheimer’s, frontotemporal and vascular dementias. Acta Neurol. Scand. 105, 355–364. https://doi.org/10.1034/j.1600-0404.2002.01147.x

Wardlaw, J.M., Allerhand, M., Doubal, F.N., Hernandez, M.V., Morris, Z., Gow, A.J., Bastin, M., Starr, J.M., Dennis, M.S., Deary, I.J., 2014. Vascular risk factors, large-artery atheroma, and brain white matter hyperintensities. Neurology 82, 1331–1338. https://doi.org/10.1212/WNL.0000000000000312

Wen, W., Sachdev, P.S., Li, J.J., Chen, X., Anstey, K.J., 2009. White matter hyperintensities in the forties: Their prevalence and topography in an epidemiological sample aged 44–48. Hum. Brain Mapp. 30, 1155–1167. https://doi.org/10.1002/hbm.20586

Woollacott, I.O.C., Bocchetta, M., Sudre, C.H., Ridha, B.H., Strand, C., Courtney, R., Ourselin, S., Cardoso, M.J., Warren, J.D., Rossor, M.N., Revesz, T., Fox, N.C., Holton, J.L., Lashley, T., Rohrer, J.D., 2018. Pathological correlates of white matter hyperintensities in a case of progranulin mutation associated frontotemporal dementia. Neurocase 24, 166–174. https://doi.org/10.1080/13554794.2018.1506039

Ylikoski, A., Erkinjuntti, T., Raininko, R., Sarna, S., Sulkava, R., Tilvis, R., 1995. White Matter Hyperintensities on MRI in the Neurologically Nondiseased Elderly. Stroke 26, 1171–1177. https://doi.org/10.1161/01.STR.26.7.1171

Zhuang, F.-J., Chen, Y., He, W.-B., Cai, Z.-Y., 2018. Prevalence of white matter hyperintensities increases with age. Neural Regen. Res. 13, 2141–2146. https://doi.org/10.4103/1673-5374.241465

